# Disturbance interacts with dispersal and niche breadth to shape scale-dependent diversity change in metacommunities

**DOI:** 10.1101/2025.03.20.644272

**Authors:** Zachary Hajian-Forooshani, Jonathan M. Chase

## Abstract

Disturbances influence the maintenance of diversity in important, but complex, ways across spatial and temporal scales. Although disturbance effects on diversity are known to be scale-dependent and taxon-specific, there is little mechanistic understanding of the processes that influence the observed context-dependency. Here, we take a theoretical approach based on metacommunity theory to examine the interaction between metacommunity processes and disturbance in shaping diversity patterns across spatial scales. We find that disturbance shapes diversity at local and regional scales in ways which can lead to either homogenization (decreases in *β*-diversity) or differentiation (increases in *β*-diversity). How it does so depend on the spatial extent of the disturbance in the landscape, the dispersal rates and niche breadth of species in the metacommunity, and whether diversity is measured immediately following disturbance or during recovery.We show that high dispersal jointly promotes the rapid recovery of local diversity and the loss of regional diversity, resulting in decreases in *β*-diversity. Niche breath buffers against diversity loss at both scales during disturbance, but interacts with dispersal to drive transient diversity loss at the regional scale after disturbance. Our results suggest that particular processes in metacommunities interact with disturbance and leave behind distinct signatures of diversity change across scales that can be used to better parse observed patterns of diversity change in empirical systems.

## I. INTRODUCTION

Disturbance can play a critical driving role for biodiversity dynamics across ecosystems (Hughes et al., 2007; Huston 2014). On the one hand, disturbance can limit the birth rates and/or increase the death rates of many species in a community, contributing to biodiversity decline (Dornelas 2010). On the other hand, many species are favored by disturbances that limit the abundance of competitively dominant species, or other strong interactors (e.g., predators) (Commander & White 2020). As a consequence, the ‘intermediate disturbance hypothesis’ and related ideas were developed to predict that the relationship between disturbance and diversity might often be hump-shaped, such as in herbaceous vegetation (Grimes 1974) and tropical forests and coral reefs (Connell 1978). The idea of a universal unimodal disturbance-diversity relationship was appealing (Fox 1979), and together with related ideas, highlighted the central roles of intraspecific competition, dispersal and non-equilibrium dynamics in structuring community assembly processes and consequent biodiversity patterns (Hubbell 1979; Wilson 1992). Like almost all patterns in biodiversity, however, the ubiquity of the unimodal disturbance-diversity relationship has been challenged both on empirical (Mackey & Currie 2001) and theoretical (Chesson & Huntly 1997, Fox 2013) grounds. As such, a general understanding of disturbance-diversity relationships remains elusive. This is likely due at least in part to the multi-scale nature of biodiversity and how it responds to disturbance (Chase 2003; Chase 2007). As a result, it will be important to simultaneously consider the complex ways that species interact endogenously within communities (competition, facilitation, predation), as well as their spatiotemporal interactions with environmental variation within a landscape, to better understand how disturbance structures the dynamics of biodiversity in space and time.

Biodiversity dynamics emerge from the interaction of local and regional scale processes (Wilson 1992; Leibold et al., 2004; Chase et al., 2019). Disturbances also occur across a range of spatial scales and can act differently on scale-specific processes within ecological communities (Turner 2010). Given the scale-dependent nature of both biodiversity dynamics and disturbances, it is perhaps unsurprising that the relationship between disturbance and diversity is often scale-dependent (Chaneton & Facelli 1991; Kaiser 2001; Hamer & Hill 2001; Chase 2003; Chase 2007; Hill and Hamer 2004; Dumbrell et al., 2008). These scale specific responses of diversity to disturbance are encompassed in the dynamics of *β*-diversity change. Whittaker’s *β*-diversity explicitly considers diversity at local (*α*) and regional (*γ*) scales simultaneously via 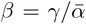 (Whittaker 1960; Whittaker 1972). An interrogation of *β*-diversity dynamics in response to disturbance allows one to understand the relative contributions of local and regional scale diversity change in shaping spatial turnover in biodiversity.

Disturbance is often associated with a reduction in *β*-diversity (Weeks et al., 2022; Roeder et al., 2018; House and Bever 2018; Tingley et al., 2016), but increases in *β*-diversity have also been observed (Socolar et al., 2016; de Carvalho et al., 2016). It is clear that both increases and decreases in *β*-diversity can emerge depending on the dynamics of both the disturbance and the community (Rolls et al., 2023). There are different kinds of disturbance that can variably influence local coexistence and species compositional variation across localities (i.e., *β*-diversity) resulting in scale-dependent patterns. For example Chaneton & Facelli (1991) found that floods decrease plant diversity at local and regional scales, but grazing increases diversity locally and reduces it at larger scales. Different ecological communities can also have contrasting responses to similar disturbances. In the face of logging in tropical forests, bird diversity can be lower at local scales, but higher across the entire landscape, while butterflies have opposite scale-dependent responses (Hill and Hammer 2004).

The emergence of taxon-specific disturbance-diversity relationships suggest the importance of varying strengths of processes within and between communities in determining their sensitivity to disturbance and their recovery. While the importance of competition in structuring the dynamics of diversity in disturbance-driven systems is well appreciated (Huston 1979; Shea et al., 2004; Miller et al., 2011; Huston 2014), dispersal and niche breath can also play a critical role. Dispersal can shape how different communities respond to disturbance (Cadotte 2007; Pedley & Dolman 2014), for example when higher dispersal rates can influence how quickly communities respond to, or recover from, disturbance (McIntyre et al., 1995; Smale 2008; Malmström 2011). While less is known about the influence of niche breadth, or the range of environmental conditions species can use, it has been shown to also be important, for example, in how spider communities respond to flooding (Lambeets et al., 2008). Disentangling the relative importance of community processes in structuring on scale-dependent responses of diversity to disturbance will help allow a mechanistic understanding of the impacts of disturbance on patterns in *β*-diversity both during and after disturbances.

Theoretical considerations of the role of disturbance on diversity have largely considered mechanisms underlying species coexistence (Huston 1979; Roxburgh et al., 2004; Miller et al., 2011; Pulsford et al., 2016). Conceptual models can help organize ideas about the interactions between disturbance and *β*-diversity (Chase 2003; Rolls et al., 2023), but these interactions have not been addressed using mathematical models. Metacommunity theory suggests that both dispersal (Mouquet & Loreau 2002; Mouquet and Loreau 2003; Matthiesen & Hillebrand 2006) and the degree to which species have specificity in habitat requirements (Pandit et al., 2002; Büchi & Vuilleumier 2014; Thompson & Gonzalez 2016) shape diversity dynamics at multiple spatial scales. It follows that these processes influence metacommunity sensitivity and response to disturbance. For example, when species have high-dispersal rates, we expect local scale *α*-diversity will recover rapidly post-disturbance due to recolonization of disturbed patches, but will likely contribute to reductions in *β*-diversity resulting in homogenization across space. Furthermore, when species have broad environmental niches, changes in regional diversity should be less sensitive to disturbances due to high occupancy at the regional scale. The resistance of diversity loss at regional scales, coupled with local extirpation makes it more likely that communities will increase in *β*-diversity post-disturbance.

Here, we explore how metacommunity processes drive changes in diversity across spatial scales in response to disturbance, with a particular focus on the impact on patterns of *β*-diversity. We base our model on previous metacommunity theory that considers spatially explicit metacommunities of competitors (e.g, Thompson et al., 2020; Guzman et al., 2021) to show how species niche breath and dispersal dynamics interact with pulse disturbances to shape diversity dynamics. We first ask how dispersal rates influence local and regional scale diversity response to disturbance. We then ask how niche breadth and dispersal interact to shape the long-term recovery patterns of disturbed metacommunities. We emphasize the importance of the relative losses in diversity between spatial scales and how non-linear diversity change at regional scales that can produce complex patterns of *β*-diversity through time. Finally, we explore how community processes differentially influence the likelihood of extinction at the regional scale during and after disturbance. Our theoretical exploration helps to make explicit expectations regarding the scale-dependent biodiversity change that can emerge following disturbance.

## II. METHODS

Our general approach is outlined in Figure 1, where we demonstrate the possible outcomes in community composition before and after a pulse disturbance and illustrate scale-dependent changes in diversity (Fig 1a.). Typical dynamics of the metacommunity are shown in Figure 1b., where the closed metacommunity is seeded onto a landscape, then the environment experiences a pulse where those environmental values are made suboptimal for the metacommunity. A range of pulse disturbance scenarios were separately simulated with different spatial extents of disturbance (Fig 1c.).

**FIG. 1.**
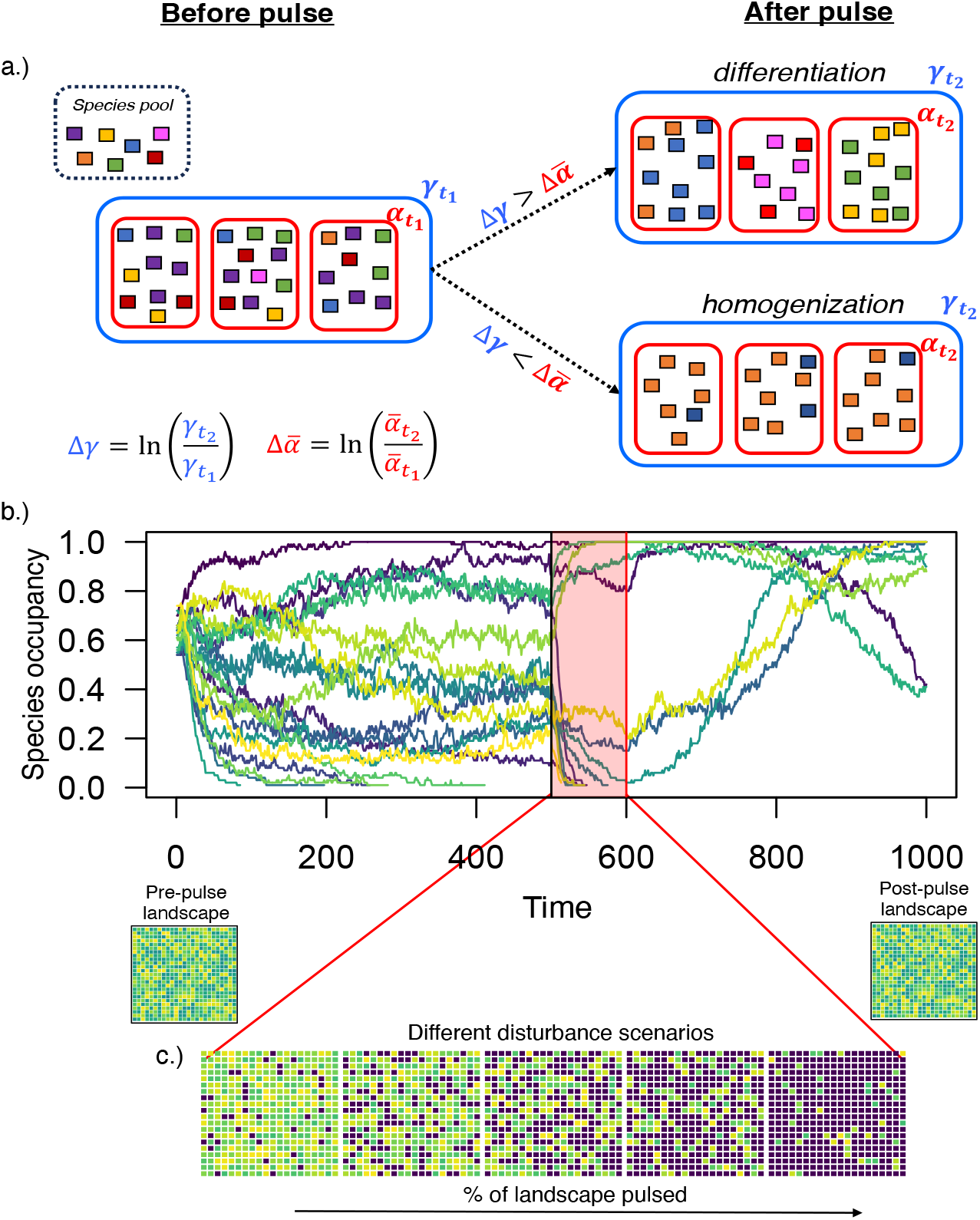
Simple pulse disturbances drive changes in diversity at local (*α*) and regional (*γ*) scales. a.) illustrates how relative the changes in diversity at different spatial scales result in compositional change (i.e. in *β*-diversity change) following disturbance. When losses at the local scale outpace losses at the regional scale, then *β*-diversity increases as patches differentiate across space. When the opposite is true and regional losses outpace local losses, then local communities become more similar to each other (i.e., homogenization) and *β*-diversity decreases. Scale specific change is calculated with log-change as shown in the figure and note that we deal with closed communities, meaning that Δ*γ* will always be less than or equal to zero. b.) Illustrates the temporal dynamics of the metacommunity simulation with pulse disturbance. Each line shows the fraction of patches occupied for a species in the metacommunity through time. At time 500, a random subset of patches in the environmental are disturbed instantaneously (vertical black line) then brought back to their initial values after 100-time steps (the vertical red rectangle). Note that species are lost during the disturbance and transient dynamics post-pulse disturbance are observed. We explore a range of disturbance spatial extents as illustrated c.), where the green/yellow squares represent habitat patches (species in metacommunity can grow optimally) and the dark purple patches represent disturbed patches (all species grow sub-optimally).

### 1. Model description

We based our model on the framework proposed by Thompson et al. (2020), modified here to incorporate environmental disturbances. Similar to other metacommunity models (e.g., Guzman et al., 2021; Wisnoski & Shoemaker; Khattar & Peres-Neto 2024), we include three key processes that structure metacommunity dynamics: density-independent growth in different environments, density-dependent competitive interactions, and dispersal in spatially explicit landscapes patches. The model takes the form of a coupled map lattice (Kaneko 1992) and its basic structure of the model is outlined in equation 1.

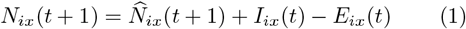

A stochastic implementation of the Beverton-Holt model that incorporates competition between populations gives the dynamics of a population in a patch (eq 2.)

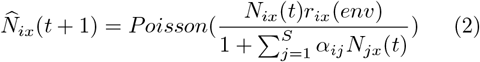

The Beverton-Holt map is the discrete time analogue to the continuous time logistic-growth model, meaning that in the absence of inter-specific competition, spatial coupling, and demographic stochasticity, that the dynamics within a patch are locally stable at a fixed point. The scaling-up of the single population Beverton-Holt map to a metacommunity results in a model that can produce relatively complicated dynamics, but the basic population dynamics of the Beverton-Holt map does not have the inherent cycling dynamics that emerge from other commonly used maps in ecology such as the Logistic-map or Ricker-map.

The growth of a population is determined by the match between the environmental condition in patch *x*, at time *t*, for species *i*, is described by *r*_*ix*_(*env*) (eq. 2).

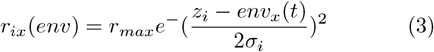

Where *env*_*x*_(*t*) represents the value of the environmental condition at time *t, z*_*i*_ the environmental optima for species *i*, and *σ*_*i*_, the niche breadth for species *i*. The term containing these parameters takes the form of a Gaussian and acts to modify the maximum growth rate of all of the species, *r*_*max*_, which is uniform across the species in the metacommunity. The niche breadth, *σ*, is a parameter whose value is shared amongst all of the species in the metacommunity for a given simulation and is the standard deviation of a Gaussian. Metacommunities with larger *σ* are those consisting of more “generalist” species that have positive growth rates in a wider range of environmental patch values, while those with smaller *σ* are more “specialist” metacommunities. Similar approaches of capturing species niche breadths with regards to growth in variable environmental conditions have been use in earlier metacommunity models (Büchi et al., 2009; Büichi & Vuilleumier 2014).

Once the growth rates of a population have been determined by their match with the local environmental conditions and the effects of both intra and inter-specific competition from co-occurring populations in the patch have been considered, demographic stochasticity is incorporated. Prior to the emigration and immigration of individuals out of a patch, the number of individuals of the population in a patch, 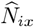, is calculated with equation 2, where the deterministic dynamics of the population become the lambda parameter of a Poisson distribution. This is an approach that simulates demographic stochasticity into the population dynamics (Shoemaker & Melbourne 2016) and necessarily relates the strength of stochasticity with the deterministic size of a population in a patch.

The spatially explicit dispersal dynamics also incorporate an element of stochasticity in the dynamics of the model. The number of species leaving a patch, *E*_*ix*_(*t*), is determined by *N*_*ix*_(*t*) draws from a binomial distribution with the probability, *a*_*i*_, which is effectively the dispersal rate of the population and approximates the fraction of *N*_*ix*_(*t*) that will leave the patch. All emigrating individuals are put into patches via *I*_*ix*_(*t*) (eq. 4), where *E*_*iy*_ are the number of individuals from patch *x* going to patch *y, M* is the total number of patches, *d*_*xy*_ is the distance between patch *x* and *y*, and *L* is the strength of exponential decrease which can be interpreted as the dispersal distance of a population

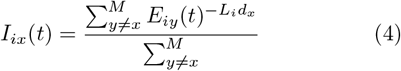

### 2. Incorporating environmental disturbance

The optimal environmental condition for species in the community, *z*_*i*_, is held constant for a simulation and is drawn from a uniform distribution *U* (*env*_*min*_, *env*_*max*_), where 0 *≤ env*_*min*_ *< env*_*max*_ *≤* 1. We constrain the values of *env*_*min*_ and *env*_*max*_ to represent a range of environmental values that are on average of higher quality for the metacommunity, something we leverage for the incorporation of environmental disturbances as described below. In our simulations we set *env*_*min*_ to 0.5, meaning that the optimal growth rates for species in the metacommunity all occur in patches with values of *env*_*x*_ *>* 0.5. This means that although there is uniform variation in species optimal environment conditions between 0.5 *−* 1, all of the optima are contained within this region. This minimum value, *env*_*min*_, represents a threshold for the metacommunity below which all species grow sub-optimally. We define disturbance when the environmental condition of a patch falls below the range for which species in the metacommunity can grow optimally (i.e. *env*_*min*_). Importantly, there remains variation in the optimal environmental conditions for species in the metacommunity, meaning that species whose optima fall closer to *env*_*min*_ will be more “disturbance tolerant” and species who optima fall closer to *env*_*max*_ will be more “disturbance sensitive”.

Although there are many ways that disturbance can be modeled, including frequency and intensity, we focus on single discrete pulse-like disturbances and how they shape scale-dependent changes in diversity. We consider disturbance to act on the environmental conditions in a patch so that its impact on species dynamics emerges indirectly through a species ability to grow in the disturbed environmental patches. This approach contrasts with theoretical studies that model disturbances that act directly on extinction rates (Büche et al., 2009) or in a density independent way (Huston 1979), but is similar to approaches where disturbance acts on species growth rates (Roxburgh et al., 2004; Miller et al., 2011).

To initialize the environmental conditions of in the model, we draw a value, *env*_*x*_, from *U* (*env*_*min*_, *env*_*max*_) for every patch in the landscape. We hold the values of *env*_*x*_ constant until the onset of disturbance at time *t*_*disturb*_, where all disturbed patches are brought down to the value *env*_*disturbed*_, where *env*_*disturbed*_ *< env*_*min*_. The duration of the disturbance lasts *t*_(*dist*−*dur*)_, where then the values of *env*_*x*_ are returned to their pre-disturbance values at time *t* = *t*_*distrub*_ + *t*_(_*dist − dur*) for the rest of the simulation. The following parameters are held constant and take the following values: *t*_(*dist*−*dur*)_ = 100, *t*_*distrub*_ = 500. To explore the extent of the landscape disturbed (fraction of patches in the environment), we randomly select a given fraction of the environment to pulse. For additional information on how we incorporate environmental disturbances in the simulation, see the supplementary material.

### 3. Quantifying scale-dependent changes in biodiversity

To quantify changes in biodiversity at different scales, we calculate the log-change in richness at local, 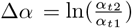, and regional, 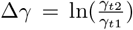, scales as outlined in Figure 1. To understand the impact of the pulse disturbance, we look at the diversity before the pulse and after the pulse. The time point prior to the disturbance, (*t*_1_), is set to, *t*_*disturb*_ *−* 1, or the time step just before the disturbance. Following the disturbance, we look at a variety of points in time for (*t*_2_) to understand both the short-term and long-term dynamics of diversity after the pulse while holding *t*_1_ constant. We also explore the dynamics of regional species loss during the pulse and after the pulse. To calculate Δ*γ* during the pulse disturbance we set *t*_2_ to the time when the disturbance stops in the landscape, while keeping *t*_1_ set to the timestep just prior to the onset of the disturbance. To calculate Δ*γ* after the pulse disturbance, we set *t*_1_ to the timestep where disturbance ends and *t*_2_ to the final timestep in the simulation.

Given the closed nature of the simulations, only negative changes are possible at the regional scale (Δ*γ*). If regional losses are greater than local losses, then spatial *β*-diversity decreases (i.e., Δ*β <* 0; if Δ*γ <* Δ*α*). If local loss outpaces regional loss, this results in differentiation across the landscape and an increase in spatial *β*-diversity (i.e., Δ*β >* 0; if Δ*γ >* Δ*α*). Finally, if changes at local and regional scales are equal, there is no change in spatial *β*-diversity (i.e., Δ*β* = 0; if Δ*γ* = Δ*α*).

### 4. Parameters of the metacommunity

We explored parameter space to determine the relevant parameter values for niche breadth and dispersal, where responses to disturbance were consistent when below the minimum and above the maximum thresholds of these dispersal and niche breadth values. For example, when dispersal rates were above d=0.01 or below d=0.00001, we find qualitatively consistent response of diversity to disturbance. Below, we highlight parameter values that shape qualitatively different diversity responses to disturbance. For example, this means we consider relatively low dispersal rates (i.e., dispersal limitation), relatively high dispersal rates (i.e., dispersal surplus), and intermediate dispersal to illustrate the qualitatively unique dynamics of diversity across the dispersal rate parameter. The same is done for the range of niche breadths. We present a wider range of parameter values in the supplementary material.

## III. RESULTS

### 1. Dispersal drives scale dependent diversity change directly after disturbance

We find that *β*-diversity in metacommunities both decrease and increase in response to pulse disturbances depending on the dispersal rates and niche breadths of the species in the metacommunity, as well as the spatial extent of the disturbance in the landscape. Dispersal limited metacommunities consistently differentiate (Δ*β >* 0) post-pulse, but the magnitude of differentiation depends on the spatial scale of the disturbance (Fig 2a). The dependence on the size of the pulse for the amount of *β*-diversity change is driven in large part by non-linear changes in diversity at the *γ*-scale. Specifically, *γ*-losses are small when a small proportion of the landscape is disturbed, but accelerate when larger proportions of the habitat are disturbed (Fig 2a). At the other extreme, when metacommunities have a surplus of dispersal, disturbance generally leads to homogenization (Δ*β <* 0), no matter its magnitude. While the non-linear losses at the *γ*-scale are qualitatively similar to the case for dispersal limited metacommunities (although with higher baseline *γ*-loss), the pattern of loss at the *α* scale is distinctly more non-linear, mirroring the *γ*-loss curve (Fig 2c). This results in largely consistent magnitudes of homogenization across the range of spatial extents of disturbance.

**FIG. 2.**
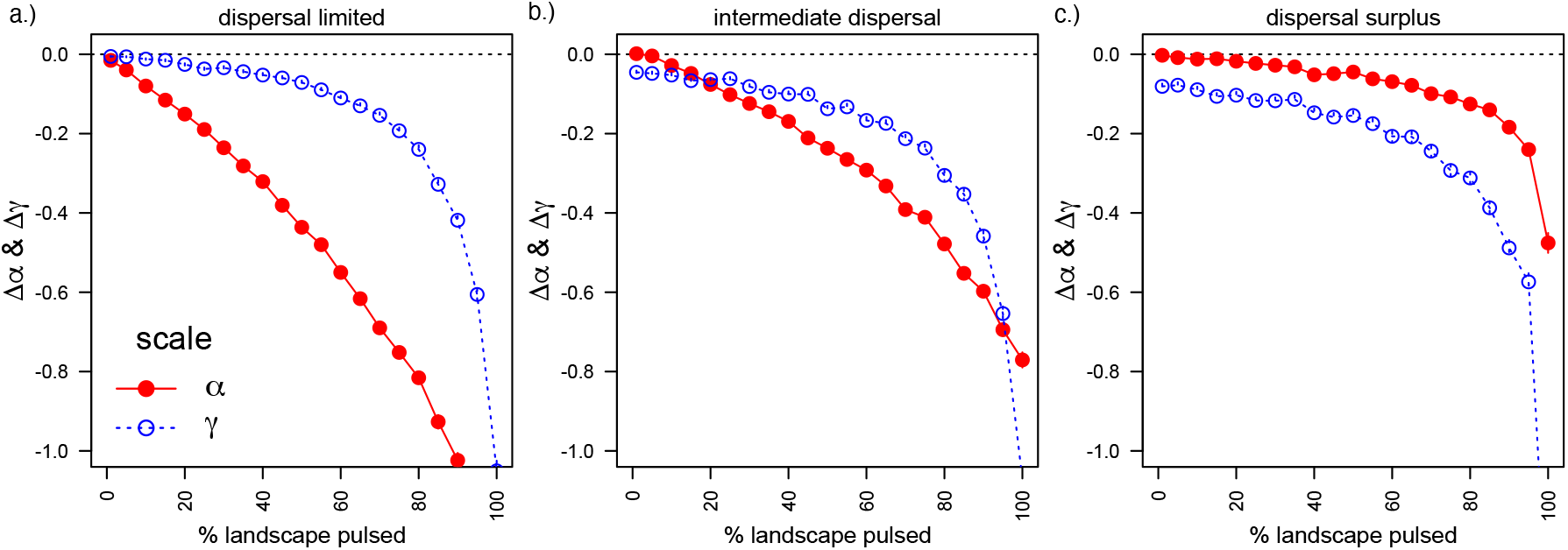
Dispersal structures the compositional shifts in metacommunity composition following pulse disturbances. a.) dispersal-limited metacommunities exhibit differentiation following pulses regardless of the extent of the pulse in the landscape, b.) metacommunities with intermediate dispersal show scale dependence in their response to pulses, and c.) dispersal-surplus metacommunities homogenize in response to pulses. Red data points show the *α*-scale changes and blue data points show the *γ*-scale changes between pre and post-pulse.

When dispersal rates are intermediate between the sur-plus and limitation scenarios, we find that the degree of homogenization and differentiation varies depending on the amount of landscape disturbed (Fig 2b). When relatively small amounts of the landscape are disturbed, losses at the *γ*-scale out-pace losses at the *α*-scale, resulting in homogenization. As the amount of landscape disturbed increases, losses at the *α*-scale begin to outpace losses at the *γ*-scale, resulting in differentiation until patterns shift again for very spatially extensive disturbances, again leading to homogenization (Figure 2b). In general, outside of the edge cases of dispersal rates (very high or very low), complex non-linear and scale-dependent patterns of loss emerge and can drive a variety of outcomes in community composition.

Although patterns can be complex, we can highlight some general effects that dispersal plays in structuring compositional shifts in biodiversity post-disturbance. When dispersal is limited, *α*-diversity will remain low after disturbance due to the lack of recolonization of patches following the disturbance (Fig 2a). This *α*-scale sensitivity of the metacommunity is the predominant driver of differentiation in the face of the disturbance and is in contrast to when dispersal rates are high. Not only do we see that *α*-scale diversity rebounds almost completely post-pulse for high dispersal metacommunities, but we also see that *γ*-scale diversity loss is relatively high even for small pulses in the landscape. This *γ*-scale sensitivity interacts with the rapid rebound in *α*-scale diversity to consistently homogenize metacommunities post disturbance (Fig 2c). By isolating the scale-dependent changes pre- and post-disturbance, we can identify the scales that contribute to compositional shifts in biodiversity patterns.

### 2. Recovery patterns of diversity are scale-dependent post disturbance

There are two components to the patterns of recovery at local and regional spatial scales which drive the *β*-diversity dynamics following disturbance. First, the recolonization of previously disturbed patches influenced the recovery of *α*-scale diversity after the disturbance. Second, when species are lost during the disturbance, the competitive dominance relationships within the community can be reorganized, resulting in regional extinctions that reduce *γ*-scale diversity through time. Below, we demonstrate how the dispersal rates and niche breadth of species metacommunities can influence these two features of metacommunity recovery. We first highlight the dynamics of scale-dependent change after disturbance, with particular emphasis on *α*-scale dynamics (Fig 3), we then show how *γ*-scale diversity loss is shaped both during and after the disturbance (Fig 4).

**FIG. 3.**
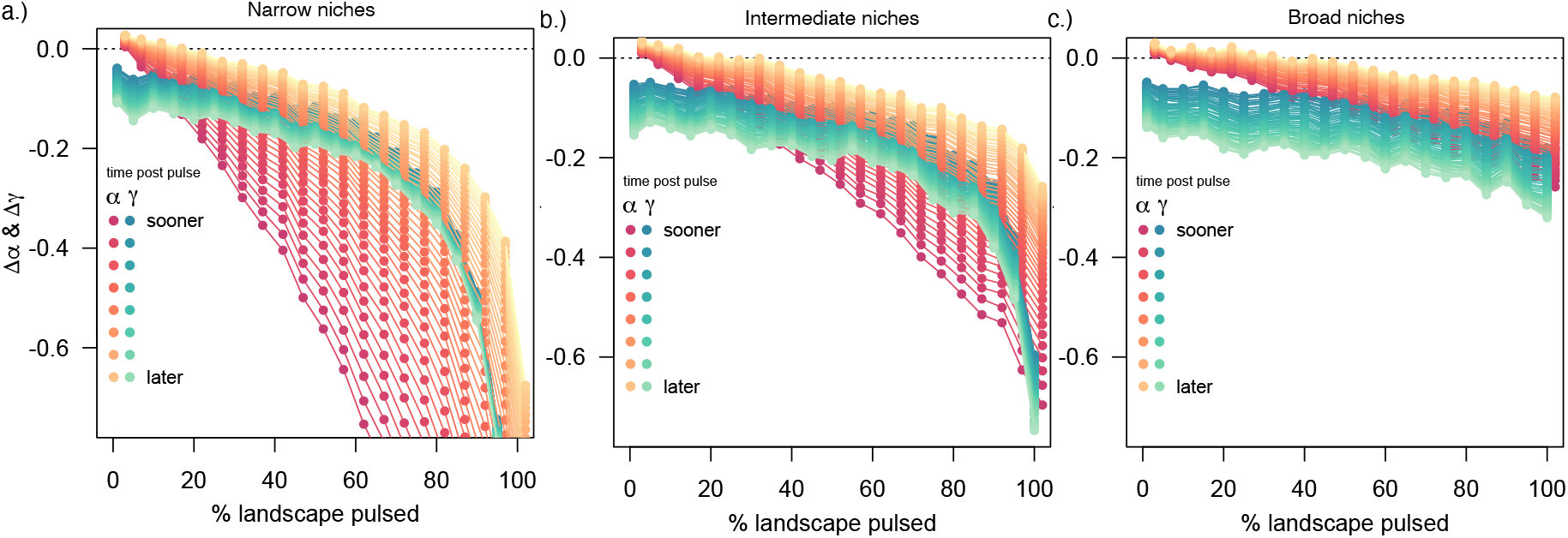
The transient scale-dependent dynamics of biodiversity post pulse disturbance for intermediate dispersal rate metacommunities with different niche breadths. Metacommunities with relatively narrow (a.), intermediate (b.), and relatively broad (c.) niches are shown. The blue lines correspond to diversity change at the *γ*-scale (regional) and the red/orange lines to the *α*-scale (local). Diversity change through time is illustrated by the colors becoming lighter. Directly after the pulse colors are dark but then they become whiter/lighter at time progresses.

**FIG. 4.**
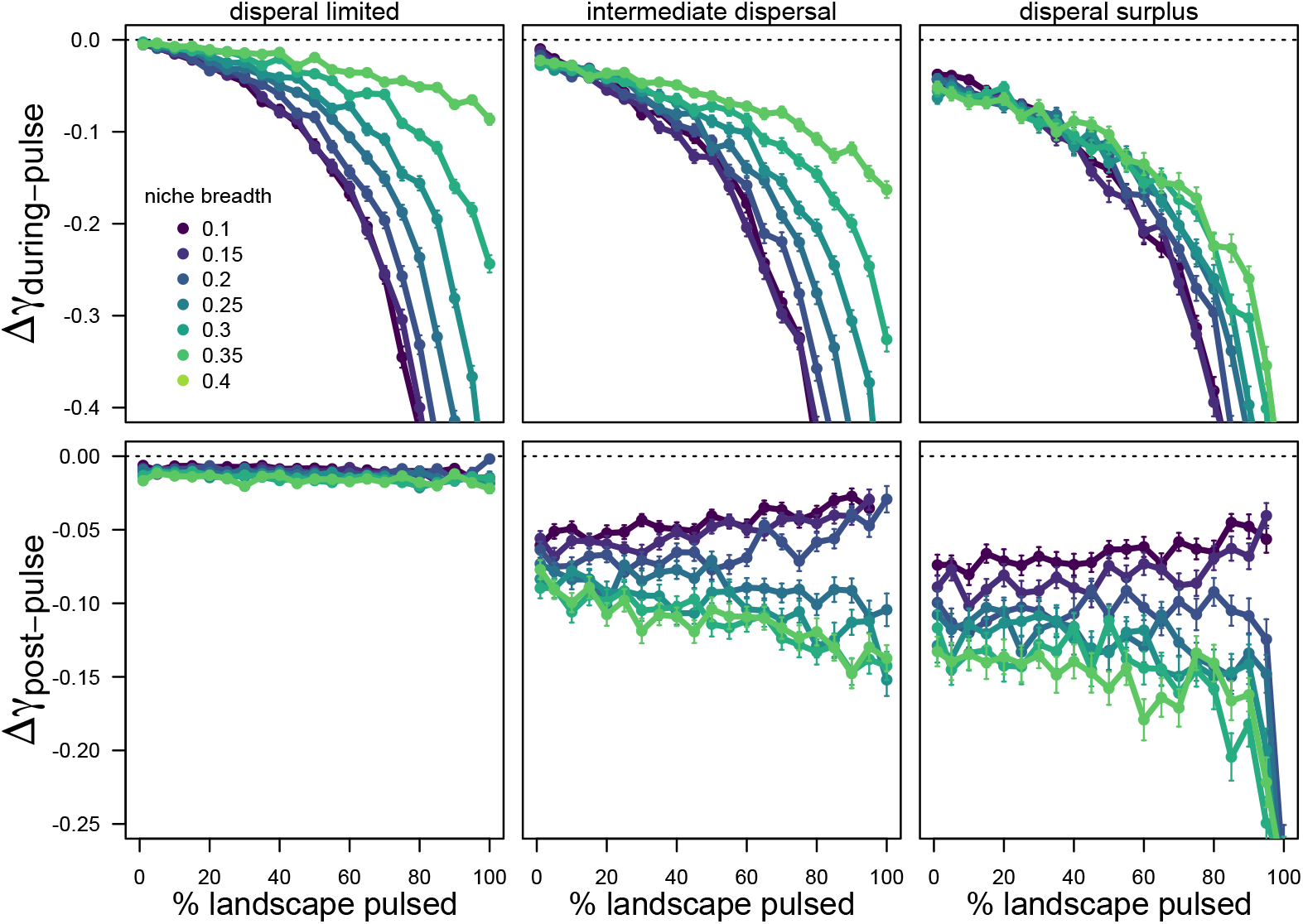
Regional diversity loss during and post disturbance structured by metacommunity niche breadth and dispersal rates. The top row shows the log-change in *γ*-diversity during the pulse and the bottom row after the pulse. The labeled columns show different dispersal rates for metacommunities from relatively low dispersal (dispersal limited) to relatively high dispersal (dispersal surplus). The multiple colors in the plot show dynamics of diversity loss for a range of different community niche breadths from more specialist (smaller values) to more generalist (larger values) metacommunities.

We find dispersal to be one of the main determinants of the metacommunity recovery process. When dispersal is limited, the metacommunity cannot recolonize patches post-pulse and the increase *β*-diversity experienced directly post-pulse (Fig 2a) remains long-term. When dispersal rates are high, the metacommunity recovers *α*-diversity almost instantaneously (Fig 2c), while driving down *β*-diversity (see Supplementary Material for meta-community recovery patterns at extremes of dispersal). Although the dynamics of recovery post-pulse for the edge cases of dispersal surplus and limitation are similar to the patterns shown in Figure 2, we find that consistent patterns in diversity dynamics at *α* and *γ*-scales post-pulse result in complex changes in *β*-diversity that are shaped by the niche breadth of the species in the metacommunity (Fig 3).

When dispersal is intermediate, both *α* and *γ*-scales tend to lose diversity during the disturbance, but they move in opposite directions through time as the community reorganizes. Furthermore, the niche breadth of species in the community strongly shapes the form of scale-dependent loss in the metacommunity (Fig 3). When niche breadth is relatively narrow, we find strong non-linear relationships at both the *α* and *γ* scales where losses accelerate as the extent of the disturbance increases (Fig 3a). This is in contrast to when niche breadths are broader, where the form of diversity loss at *α* and *γ* scales are approximately linear (Fig 3c). At intermediate niche breadth, *α*-scale diversity decreases in an approximately linear manner with the extent of disturbance, while-scale losses are non-linear (Fig 3b). The intersection of losses at these two spatial scales results in scale-dependent patterns of *β*-diversity following disturbance. This highlights how the expected changes of *β*-diversity following a disturbance can depend on both the spatial extent of the disturbance and how recently it has taken place.

### 3. Regional extinctions during and after disturbance respond differentially to community processes

After disturbance, recolonization of disturbed patches drive increases in *α*-diversity, but gradual declines in *γ*-diversity through time. This is due to the fact that species go regionally extinct during the disturbance, resulting in a novel subset of the species pool with subsequently reshaped competitive dominance. We find that regional extinctions during and after disturbance are shaped by the interaction of niche breadth and dispersal in the metacommunity (Fig 4). Dispersal-limited meta-communities lose *γ*-scale diversity only during the disturbance, and have no subsequent transient species loss after (Fig 4 – dispersal limited). These losses are determined by niche breadth, where generalist metacommunities are less sensitive to disturbance than specialists. On the other hand, dispersal-surplus metacommunities show qualitatively different patterns with respect to the importance niche breadth during and after the disturbance. Being generalist or specialist has little impact on *γ*-diversity loss during disturbance, but generalist meta-communities experience much more regional extinction after disturbances (Fig 4 – dispersal surplus). In general, we find that change in *γ*-scale diversity is more sensitive to the spatial extent of the disturbance during the disturbance than after, with strong non-linear losses in diversity during the pulse and weak relationships post-pulse (Fig 4).

## IV. DISCUSSION

In all, we show that environmental disturbances can lead to both homogenization (Δ*β*-divesrity *<* 0) and differentiation (Δ*β*-diversity *>* 0) depending on the nature of the disturbance, the characteristics of the species in the metacommunity, and the time in which diversity is measured. Our results emphasize the role of dispersal and niche breadth of the species in a metacommunity in shaping *α*-scale (local) and *γ*-scale (regional) diversity changes and how the relative changes of diversity at these two scales determine the strength and direction of *β*-diversity change following disturbance. Furthermore, we identify how these processes in metacommunities interact to make diversity more or less sensitive to losses at different spatial scales, as well as how scale-dependent diversity patterns recover through time. Although the interplay of dispersal dynamics and niche breadth can generate a complex array of spatiotemporal outcomes in the face of simple disturbance, we suggest that the scale-dependent interrogation of changes in biodiversity can provide intuition that may help understand how different communities will respond to disturbance in variety of contexts.

The fact that biodiversity exhibits a wide range of responses to environmental changes is well appreciated (Vellend 2010), even when considering a single driver such as disturbance (Mackey & Currie 2001; Burkle et al., 2015; Ribeiro-Neto et al, 2016; Catano et al., 2017). We find that even in simplified scenarios, changes in both the magnitude and direction in *β*-diversity can emerge from the same metacommunities experiencing disturbances with different spatial extents (Fig 2b & Fig 3). Similar relationships between disturbance size/intensity and changes in *β*-diversity are sometimes thought to operate through the generation of environmental heterogeneity, where increases in heterogeneity emerge from isolated disturbances and serve to differentiate communities, while large disturbances can homogenize environmental conditions (Burkle et al., 2015; Limberger & Wickham 2012; Östman et al., 2006). When environmental filtering is strong, we also expect that biodiversity will respond in concordance with disturbance induced changes environmental heterogeneity (Barrett et al., 2023). Changes in *β*-diversity have been shown empirically to be determined by the scale of disturbances, for example observations of forests in response to fire (Burkle et al., 2015; Weeks et al., 2022), and in laboratory experimental metacommunities of protists, where local disturbances increase *β*-diversity and regional disturbance decreases *β*-diversity (Limberger & Wickham 2012).

There is a rich theoretical literature and long history of exploring the relationships between disturbance and diversity, but notably few address the response of diversity across spatial scales and have instead focused on the mechanisms underlying local coexistence (e.g., Roxburgh et al., 2004). As we show here, research that explicitly considers diversity dynamics at multiple spatial scales can help provide more mechanistic understanding of community sensitivity to, and recovery from, disturbance (Whitlatch et al., 1998; Jaquet et al., 2022). For example, Tatsumi et al., (2020) analyzed changes in diversity across scales through the lens of colonization and extinction to understand the drivers of *β*-diversity changes following disturbance. Additionally, Cunillera-Montcusí et al., (2021), highlighted the central role of dispersal distance in shaping patterns of resilience and recovery to disturbance. The joint consideration of scale-explicit diversity dynamics and they are related processes within metacommunities, can help build a more mechanistic understanding of how environmental disturbance influences biodiversity dynamics.

Although there are a number of features of real ecosystems that are not considered in our simplified approach here, qualitatively similar patterns of biodiversity change produced by our model have been reported empirically. For example, a meta-analysis by Catano et al., (2017) show that low dispersal rates and dispersal limitation tends to increase *β*-diversity while high dispersal decreases *β*-diversity (Fig 2 & Fig S4). The sensitivity of specialist species (narrower niches) to disturbance that we expect (Fig 3 & Fig 4) has also been reported in a variety of different systems, including birds (Devictor et al., 2008) and butterflies (Kitahara & Fujii 1994; Kitahara et al., 2000). The relative importance and interaction between niche breadth and dispersal in response to disturbance is less well understood and likely community and disturbance dependent (Lambeets et al., 2008). For example, Mabry and Fraterrigo (2008) find that habitat specialization is a more important determinant of community structure in response to disturbance than dispersal strategy in North American deciduous forests. Our results suggest strong interactions between these two processes within metacommunities, but also further highlight the importance competitive interactions.

We found that competition plays a role in the transient reorganization of the metacommunity following a disturbance due to the regional loss of species during the pulse (Fig 4). It is well appreciated that the loss of a species can cascade through the community (Allesina & Bodini 2004; Petchy et al., 2008). There are a number of ways that competitive networks can be reshaped by extinction that will cause novel extinctions in the community post-disturbance. For example, competitive release from a species lost during disturbance can lead to new outcomes that result competitive exclusion (Fowler 2010), or the loss of species that take part in intransitive structures in the network can also result in competitive exclusion (Laird & Schamp 2006). Although competition plays an important role in our observed results, an explicit interrogation of the competitive matrix is outside the scope of our current study. We instead took the commonly-used approach of using random matrices of competitive interactions to understand some general expectations (May 1972; Allesina & Tang 2015; Akjouj et al., 2024). Exploring how the structure of competitive networks influences the sensitivity of communities to disturbance and shapes post-disturbance recovery dynamics is largely an outstanding question. Potentially important considerations include the trade-offs that exist in some communities between competitive ability and other species traits, such as dispersal (Tilman 1994) and disturbance tolerance (Violle et al., 2010). While some trade-offs associated with competition have may be important in shaping metacommunity responses to disturbance, it is not entirely clear which trade-offs might be most important (Seifan et al., 2013) and when (Haddad et al., 2008).

While it is clear that biodiversity is changing around the globe, there is considerable variation in the observed trends through time and across space (Dornales et al. 2015, Vellend et al. 2014, Leung et al. 2019; Blowes et al. 2024; van Klink et al., 2024). Building a theory-informed intuition regarding the mechanisms that shape the observed dynamics of biodiversity remains an important challenge (O’Sullivan et al., 2021; Storch 2022; Okie & Storch, 2025). Here we work towards such an intuition by exploring how processes in metacommunities shape their responses to disturbance. We show how scale-dependent changes in diversity can be understood through the interaction of dispersal and niche breadth in metacommunities, which leave behind distinct signatures on diversity change during and after disturbance. Our approach provides a framework for mechanistic predictions regarding how metacommunities that are differentially structured by these processes should respond to disturbance. Building connections between functional approaches that can quantity the importance of these processes in metacommunities will likely go a long way to building predictive frameworks for how biodiversity will response to anthropogenic drivers.

## ACKNOWLEDGMENTS

Z H-F would like to thank Iris Saraeny Rivera Salinas, Kristel Sanchez, Chatura Vaidya, Otso Ovaskainen, Ferdinand LaMothe and the Biodiversity Synthesis group at iDiv for feedback and discussion at various stages of the project.

## VI. SUPPLEMENTARY MATERIAL

### 1. Incorporating environmental disturbance in the metacommunity model

We take an approach that conceptualizes disturbance as acting on the environment in such a way that it reduces the growth rates of species in the metacommunity. We assume that there is variation in species optimal growth rates but that all growth rates occur above a threshold (the metacommunity threshold). Values of the environment in patches can be thought of as representing some characteristic of the environment, such as moisture, and we assume that species will have differences in optimal performance along the moisture gradient but that there exists some threshold across which no species will perform optimally (Fig S1b). We set all patches in the environment to start above this threshold and to simulate disturbance, we take some fraction of the patches in the environment (60% in Fig S1) and bring them below this threshold (to the environmental value of 0.1 for all simulations here) resulting in suboptimal growth of all species in the metacommunity in the effected patches. An example of the temporal dynamics of patch disturbance is shown in Figure S1, where we have 100 patches and 60% of them starting at t=500 until t=600. Note that each patch is initialized with a random environmental value that falls within the range optimal growth values for all species and remains static until the pulse. Furthermore, the pulse disturbance starts and stops on a single timestep and that post-pulse the environmental patches return to their initial values (Fig S1a).

**Figure S1.**
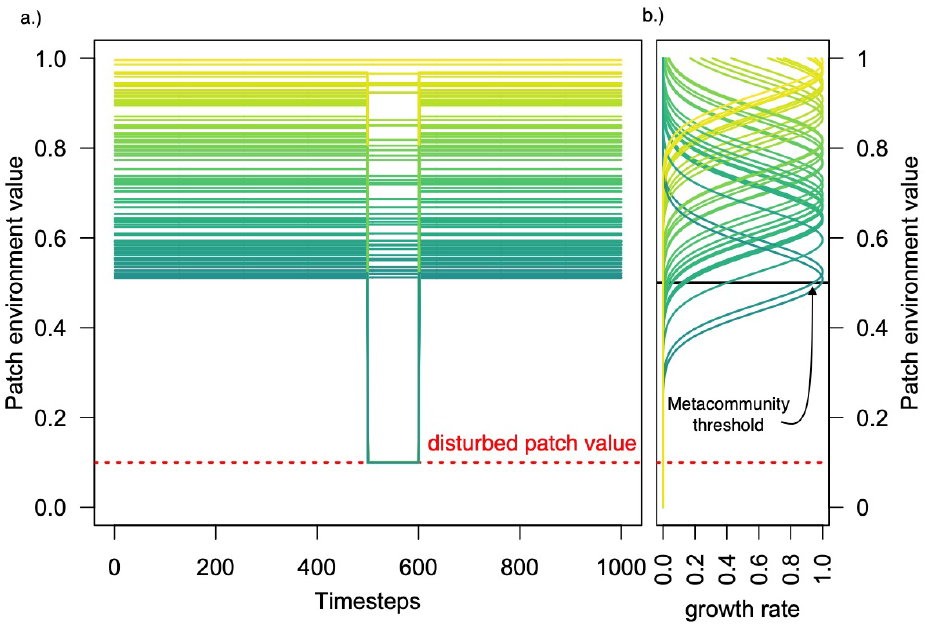
The dynamics of the environmental disturbance in the simulations. a.) the environmental values of each patch in a landscape with 100 patches, where a line represents a patch and the color where it falls on the environmental patch value gradient. At time 500 the disturbance is initiated on 60% of the patches which are brought down to a disturbed value where all species will grow sub-optimally in the metacommunity. b.) illustrates the distribution of environmental growth curves for the metacommunity. The metacommunity threshold is denoted which is a threshold below which no species grow optimally in the metacommunity. Note that species closer to this metacommunity threshold will be more disturbance tolerant that species far from it.

**Figure S2.**
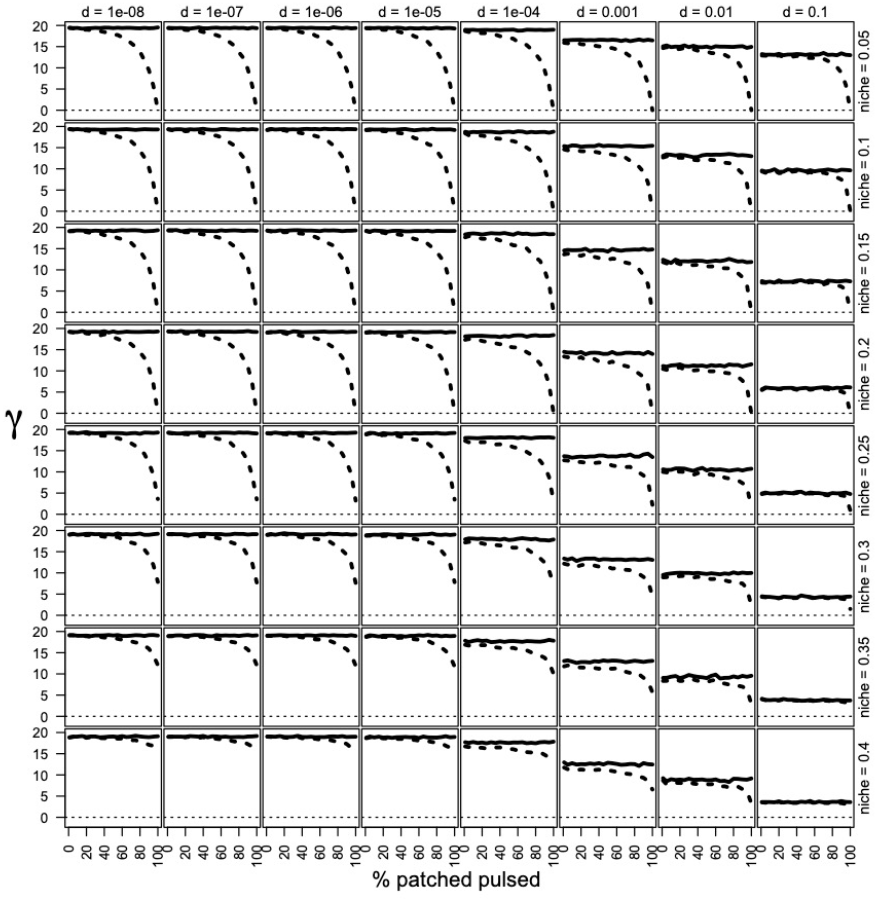
The values of *γ* diversity across a range of metacommunity parameters. The columns show the dispersal rates and the rows illustrate the niche breadths. The solid line is the *γ* diversity prior to the pulse and the dashed line after the pulse.

**Figure S3.**
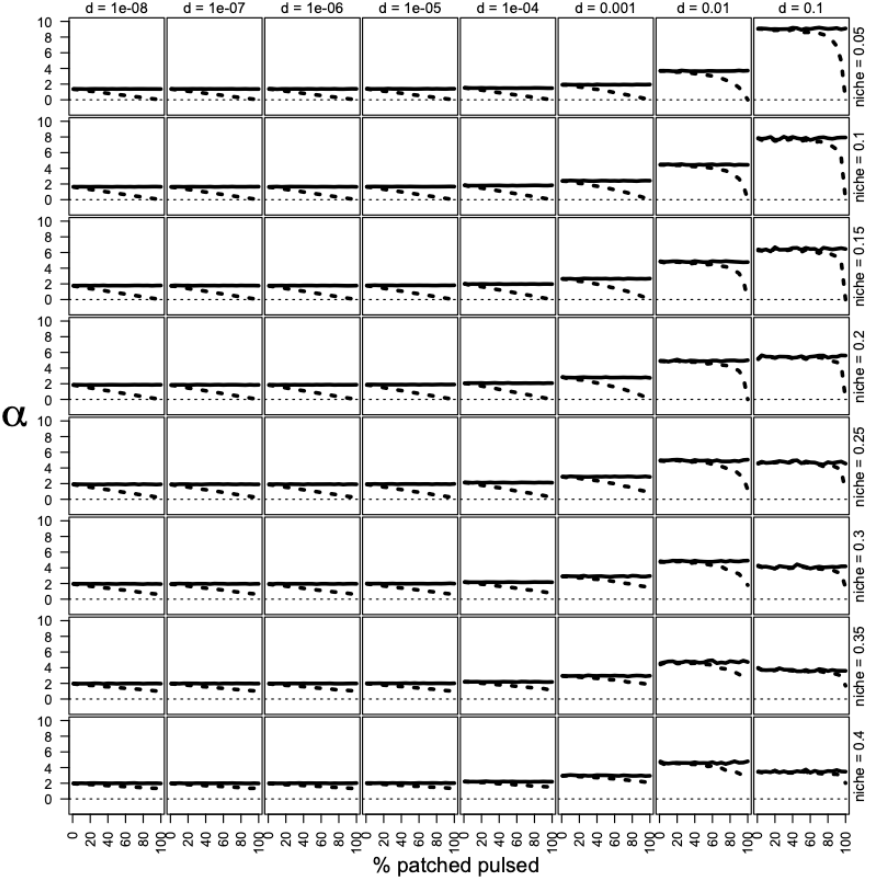
The values of *α* diversity across a range of metacommunity parameters. The columns show the dispersal rates and the rows illustrate the niche breadths. The solid line is the *α* diversity prior to the pulse and the dashed line after the pulse.

### 2. Effects of dispersal and niche breadth on α and γ-diversity

Here we illustrate the joint effects of niche breadth and dispersal on *α* and *γ*-scale diversity in the metacommunity model. At the regional scale, we see an interactive negative effect of both increase dispersal rate and increasing niche breadth on *γ* diversity (Fig S2). As we increase dispersal rates, we see declines in the *γ* richness of the metacommunity, and a similar effect of declining *γ*-diversity is observed as we increase the breath of the niches in the metacommunity. At the *α* scale, we see that increasing dispersal has a positive effect on diversity for narrow niche breadths but a hump shaped effect when niche breadth is broader (Fig S3). These negative effects can be understood and emerge from competition between species in the metacommunity. As dispersal increases, species can move through the landscape and competitively exclude other species reducing diversity. Furthermore, as the overlap in environment requirements for species increases with increasing niche breadth, competitive exclusion will also occur more frequently. In the main body of the manuscript, we used values of dispersal and niche breadth that illustrate the qualitatively distinct outcomes we observed in the parameter space of the model.

**Figure S4.**
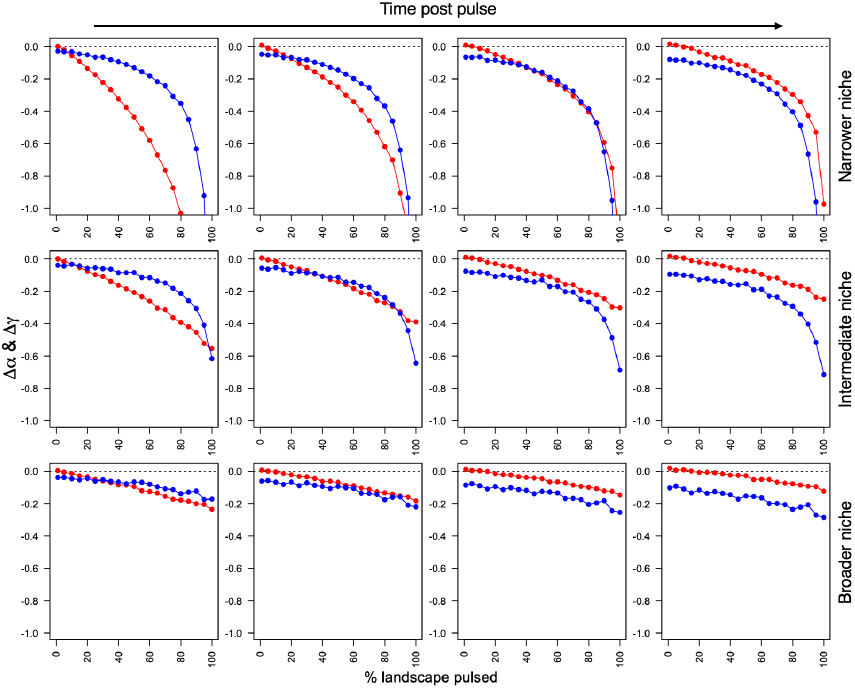
The transient dynamics of recovery for metacommunities post pulse. The parameters used here are the same as the parameters shown in the main body of the manuscript for Figure 3. (i.e., intermediate dispersal). The left most panels show the scale-dependent diversity (*γ* in blue and *α* in red) just after disturbance. As panels move to the right the recovery of the metacommunity proceeds in time.

### 3. Scale-dependent transience shaped by dispersal and niche breadth

The dynamics of scale-dependent transience in diversity change post-pulse is strongly structured by the both the niche breadth and dispersal of the metacommunity. The main body of the manuscript highlights the ability of niche breadth to create transient patterns of differentiation and homogenization post pulse for intermediate dispersal values in metacommunities. The temporal dynamics of recovery can be seen for the same values of dispersal in the manuscript but plotted out at different snapshots (Fig S4).

Here we build on the results from the main body of the manuscript and highlight the how dispersal limitation and dispersal surplus structure changes at both *α* and *γ*-scales. When dispersal is limited the change in both *α*-diversity and *γ*-diversity is minimal which results in consistent differentiation though time (Fig S4a). Metacommunities that have dispersal surplus see *α*-scale diversity rebound towards to pre-pulse states relatively rapidly (Fig S4c). For narrow and intermediate niches, we see the that differentiation can occur directly after the pulse but quickly moves towards homogenization as time progresses.

**Figure S5.**
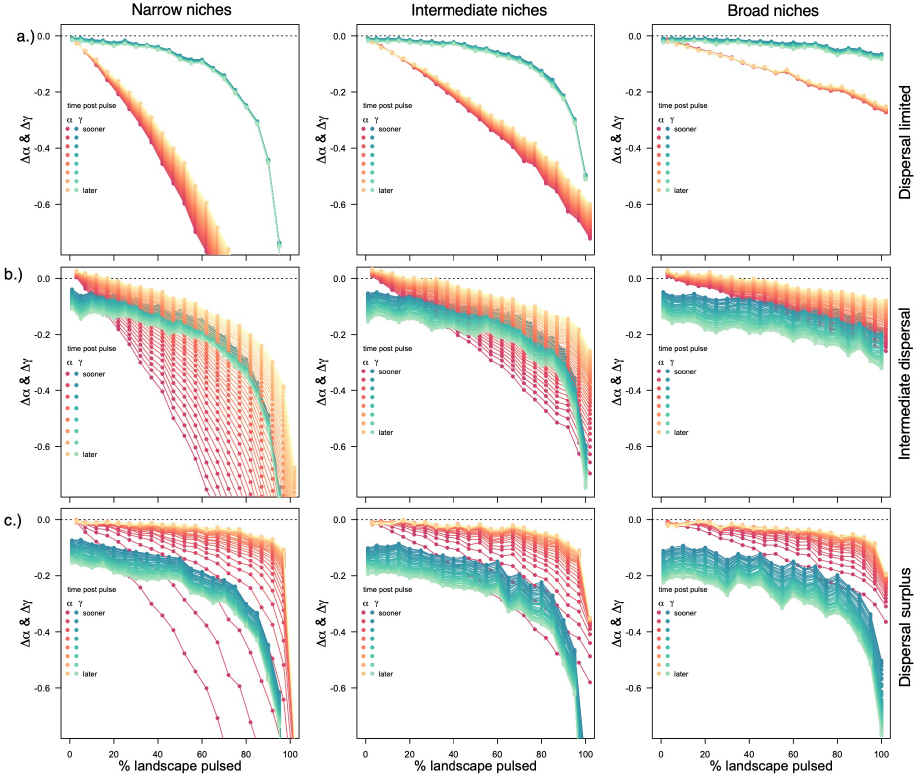
The transient dynamics of metacommunities at different points in time post-pulse. b.) is reproduced here but seen in the main text of the manuscript. a.) shows dispersal limited metacommunities, and c.) dispersal surplus metacommunities. The blue lines show *γ*-scale diversity and red lines show *α*-scale diversity. As the lines become lighter they show the values of change as time progresses (darker right after pulse and whiter later after pulse).

